# Sexual behaviour in a murine model of Mucopolysaccharidosis type I (MPS I)

**DOI:** 10.1101/705913

**Authors:** Ana Beatriz Barbosa Mendes, Cinthia Castro do Nascimento, Vânia D’ Almeida

## Abstract

Mucopolysaccharidosis Type I (MPS I) is a rare genetic lysosomal storage disease caused by a mutation of *IDUA* gene. *IDUA* codes for α-L-iduronidase (IDUA), a lysosomal hydrolase that degrades glycosaminoglycans (GAGs): heparan sulphate and dermatan sulphate. GAGs are structural and signalling molecules that have a crucial role in controlling a variety of cell functions and their interaction with extracellular matrix. Because of GAG’s widespread action in cellular metabolism, MPS I is a progressive and disabling multisystemic disorder. Nowadays, the therapies availability allowed patients to reach the adult life and the consequences of the disease in their reproductive system is still little known. We aimed to investigate whether IDUA disruption influences sexual behaviour and sexual steroid production in male and female MPS I mice. We used 3 and 6-month-old male and 3-month-old female *Idua*+/_ and *Idua*−/− mice to evaluate typical rodent copulatory behaviours. In males we observed the frequency and latency of mounts, intromissions and ejaculations. In females we evaluated the lordosis quotient. We also analysed the locomotor capacity of mice in the open field test, since copulatory behaviour requires mobility. We also quantified steroidal hormonal levels in plasmatic samples. We detected an increase in the latencies of intromissions in male copulatory behaviour of *Idua*−/− males when compared to *Idua*+/_. However, the number of intromissions was not statistically different between groups. No parameter of female sexual behaviour was statistically different between control and knockout females. In both sexes, we detected diminished mobility in *Idua*−/− mice. Plasma hormone levels did not differ between *Idua*+/_ and *Idua*−/− mice, both in males and females. We concluded that in the considered time point of MPS I progression, mice are able to perform sexual behaviour, but the male performance may be influenced due to the motor disability predicted to MPS I.

## Introduction

Mucopolysaccharidosis Type I (MPS I) is a lysosomal storage disorder caused by a disfunction of α-L-iduronidase (IDUA), a lysosomal hydrolase coded by the *IDUA* gene, responsible for degrading two specific glycosaminoglycans (GAGs): heparan sulphate (HS) and dermatan sulphate (DS). IDUA deficiency results in the storage of both GAGs in the intra and extracellular compartments, leading to disturbances in cell signalling, lysosomal traffic and tissue architecture [1–2].

Many cell types, tissues and organs are affected by IDUA deficiency, making MPS I a systemic disease [1]. Clinical signs, such as hepatosplenomegaly, impaired growth, dysostosis multiplex and macroglossia are easily detected even in more attenuated phenotypes [3]. A wide phenotypical spectrum is described for MPS I, from attenuated to severe forms [1].

With the advancement of therapeutic strategies, such as enzyme replacement therapy and bone marrow transplantation, there has been an increase in both quality of life and life expectancy of MPS I patients [4]. Increasingly efficient therapies have enabled patients to reach puberty and, in some cases, conceive and bear children [5–7].

However, the effects of MPS I on reproduction are mostly unknown. The little research available suggests that GAG deposition may influence morphological and physiological reproductive parameters in MPS I [8]. Our group has observed significant morphological changes in testis, sperm and epididymis of MPS I male mice [9–10]. Moreover, signs of precocious puberty in MPS boys have been reported [11], indicating it may also affect hormonal production and function.

Besides morphological changes, cognitive impairment can be observed in severe phenotypes of MPSs in general [12–13]. Impaired reproductive performance was detected in MPS VII model, which was attributed to the impaired mobility and cognitive function of untreated male and female knockout models [12].

Thus, the aim of the present study was to evaluate the effect of IDUA deficiency in male and female copulatory behaviours, as well as in plasma steroid hormone concentration in a MPS I murine model.

## Materials and Methods

### Animals

*Idua* −/− (KO) model, developed by Ohmi and colleagues [14], was generated by targeted disruption of the *Idua* gene. We maintained the mice colony at our institutional animal facility (Department of Psychobiology of Universidade Federal de São Paulo) through mating between heterozygotes. Mice were housed in temperature-controlled rooms, with a light/dark cycle of 12 hours, and free access of water and food. Genotyping was performed by polymerase chain reaction, with the following primers: forward: 5’-CAG ACT TGG AAT GAA CCA GAC-3’; reverse 1: 5’-GTT CTT CTG AGG GGA TCG G-3’ and reverse 2: 5’-ATA GGG GTA TCC TTG AAC TC-3’, as previously described [13].

### Experimental design

All adopted procedures were in accordance with the Ethical Research Committee from Universidade Federal de São Paulo (CEUA # 4226130214 and 7148250117). We used 30 animals: 10 female (2 *Idua*+/−; 3 *Idua*+/+ and 5 *Idua−/−*) and 11 male (3 *Idua*+/+; 2 *Idua*+/− and 6 *Idua*−/−). Since MPS I is a recessive genetic disorder and heterozygotes manifest normal phenotype with no pathological behaviour or morphological characteristics, dominant homozygous and heterozygous mice were used to constitute the control group.

### Male sexual behaviour

We evaluated male sexual behaviour in two different time points of disease progression: 3 and 6 months of age. Fifteen days before testing, each male mouse was housed with an experienced *Idua*+/− female, for one week, to acquire sexual experience. Males spent 24 hours isolated to minimise mounting from their cagemates before being returned to their cage, where they remained abstinent for the following week.

Copulatory behaviour was observed for 30 minutes during the dark phase, under dim red lighting. We recorded and registered the number of three typical rodent copulatory behaviours: mounts, intromissions and ejaculations, as well as the latencies for each behaviour (Adapted from Kikusui et al., 2013) [15].

### Female sexual behaviour

Since female required daily monitoring of the oestrous cycle, we decided to evaluate them only at three months of age to minimize stress. Two-month-old female mice had their oestrous cycle registered for three complete cycles by vaginal lavage. When the third proestrus phase was recorded, the mice were submitted to sexual behaviour testing. The test was also performed during the dark phase, under dim red lighting, and videotaped for later scoring. Prior to the test, a healthy and sexually experienced male mouse was habituated to a neutral cage for five minutes. Following habituation, the subject was placed in the cage and they were observed for 15 minutes or until the completion of 20 mounts. This analysis was repeated for the two following proestrus phases, detected also by vaginal lavage [16]. The lordosis behaviour was quantified and normalised by the number of mounts performed by the male mice through calculation of the lordosis quotient (LQ = [lordosis frequency/mounts] *100) [16].

### Open field test

Locomotor and exploratory activities were assessed one day before male and female sexual behavioural testing, to verify whether motor impairment could interfere on sexual performance. The test was conducted in a circular arena comprised of 19 quadrants of equal area (7 central and 12 peripheral). Mice were placed in the centre of the arena and their activity was recorded for five minutes (adapted from Chinen et al., 2006) [17]. We registered the number of crossings of the quadrant line with all paws, the percentage of peripheral and central crossed quadrants, the frequency of rearing, which indicates exploratory behaviour, and the frequency of grooming, which indicates anxious-like behaviour, as a control measure.

### Hormonal plasmatic measurements

Following behavioural testing, 3-month-old female and 6-month-old male mice were euthanized by decapitation. Blood was collected in heparinised tubes, homogenised and centrifuged at 3000 *g* for 10 minutes. Plasma was collected and stored at −80° C until dosage. Progesterone, testosterone and 17-β-oestradiol plasmatic levels were determined by ELISA (Enzo Life Sciences).

### Statistical analyses

Data were analysed in SPSS. We used generalised linear models or generalised estimating equations, adjusted for the data distribution as necessary. Significance was considered as p<0.05.

## Results

### Open field tests

#### Males

KO mice had lower horizontal activity (number of quadrants crossed) when compared to control group (χ^2^ = 7.31, p = 0.007), independently of age (age: χ^2^ = 1.32, p = 0.25; interaction: χ^2^ = 0.70, p = 0.79) (Fig 1A). Interestingly, KO mice crossed more central quadrants than *Idua*+/_ (χ^2^ = 8.08, p = 0.004), also independently of age (age: χ^2^ = 0.18, p = 0.67; interaction: χ^2^ = 0.20, p = 0.89) (Fig 1B). No effect of genotype (χ^2^ =0.64, p =0.42), age (χ^2^ = 1.31, p =0.25), or their interaction (χ^2^ =0.001, p =0.98) was found on episodes of grooming (Fig 1C). Vertical activity, indicated by the number of rearing, was decreased in KO mice compared to controls (χ^2^ = 6.67, p = 0.01), independently of age (age: χ^2^ = 0.019, p = 0.89; interaction: χ^2^ = 0.54, p =0.46) (Fig 3D).

**Figure 1:**
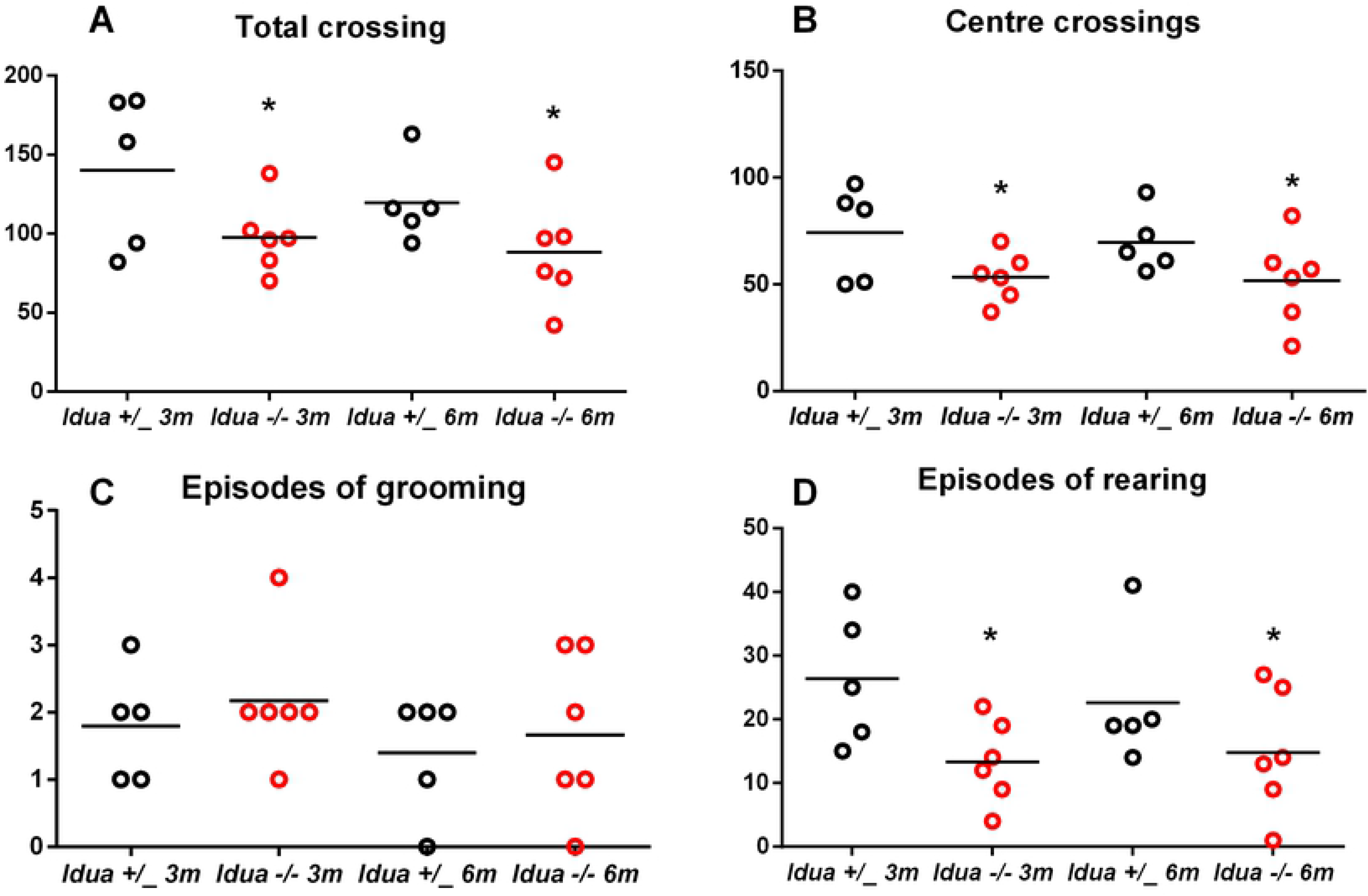
Analysis of male locomotor activity. Three and 6 months old male *Idua*+/_ and *Idua*−/− were submitted to open filed test during 5 minutes in the previous day of sexual behaviour test. The number of central and peripheral quadrants was counted, as well as the number of episodes of rearing (vertical activity) and grooming. Statistical analysis: generalized estimating equations. *p<0.05 compared to control group.

#### Females

KO females presented lower general mobility in the open field test. They travelled through fewer total (χ^2^ = 4.65, p =0.031) and central (χ^2^ = 7.62, p =0.006) quadrants when compared to *Idua*+/_ (Fig 2A-B). No difference was found in grooming (χ^2^ = 2.62, p = 0.105) (Fig 2C) or rearing (χ^2^ = 2.91, p = 0.088) (Fig 2D) frequency.

**Figure 2:**
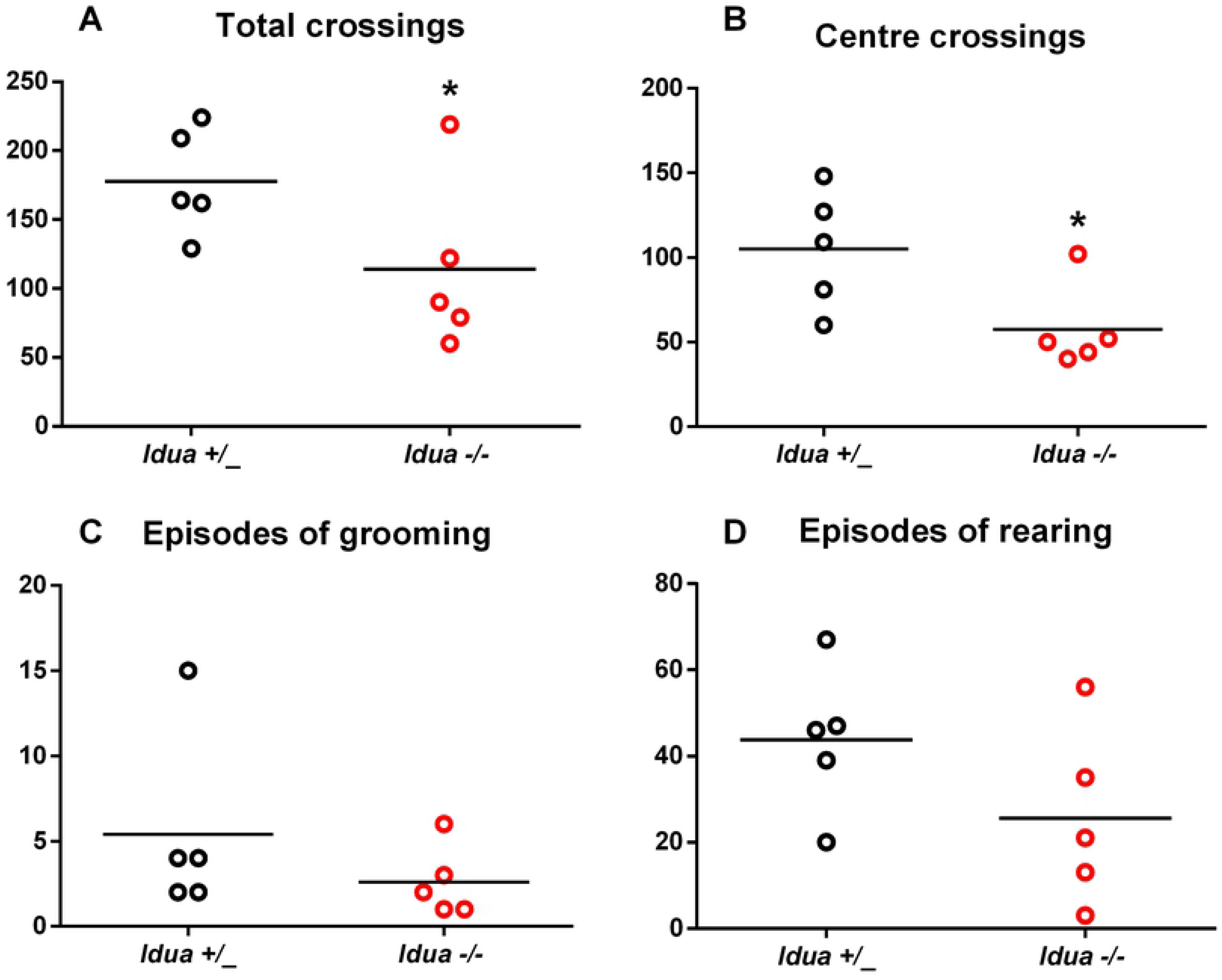
Analysis of female locomotor activity. Three-month-old female *Idua*+/_ and *Idua*−/− were submitted to open field test for 5 minutes the day before sexual behaviour testing. The number of central and peripheral quadrants was counted, as well as the frequency of rearing (vertical activity) and grooming. Statistical analysis: generalized linear models. *p<0.05 compared to control group.

### Male sexual behaviour

No effect of age (Mounts: χ^2^ = 1.09, p =0.30; intromissions: χ^2^ =0.23, p =0.64), genotype (Mounts: χ^2^ = 0.048, p =0.83; intromissions: χ^2^ = 1.55, p =0.21) or their interaction (Mounts: χ^2^ = 0.97, p =0.32; intromissions: χ^2^ = 0.012, p =0.91) was found on number of mounts and intromissions (Fig 3A-C). However, KO mice presented greater latencies of mounts (χ^2^ = 18.79, p <0.001) and of intromissions(χ^2^ = 9.71, p =0.002) when compared to wild types, independently of age (Mounts: χ^2^ = 1.45, p =0.23; intromissions: χ^2^ = 2.95, p = 0.086). No effect of the interaction between age and genotype was found on either mount (χ^2^ = 2.93, p = 0.087) or intromission latencies (χ^2^ = 1.80, p = 0.18) (Fig 3D-F).

**Figure 3:**
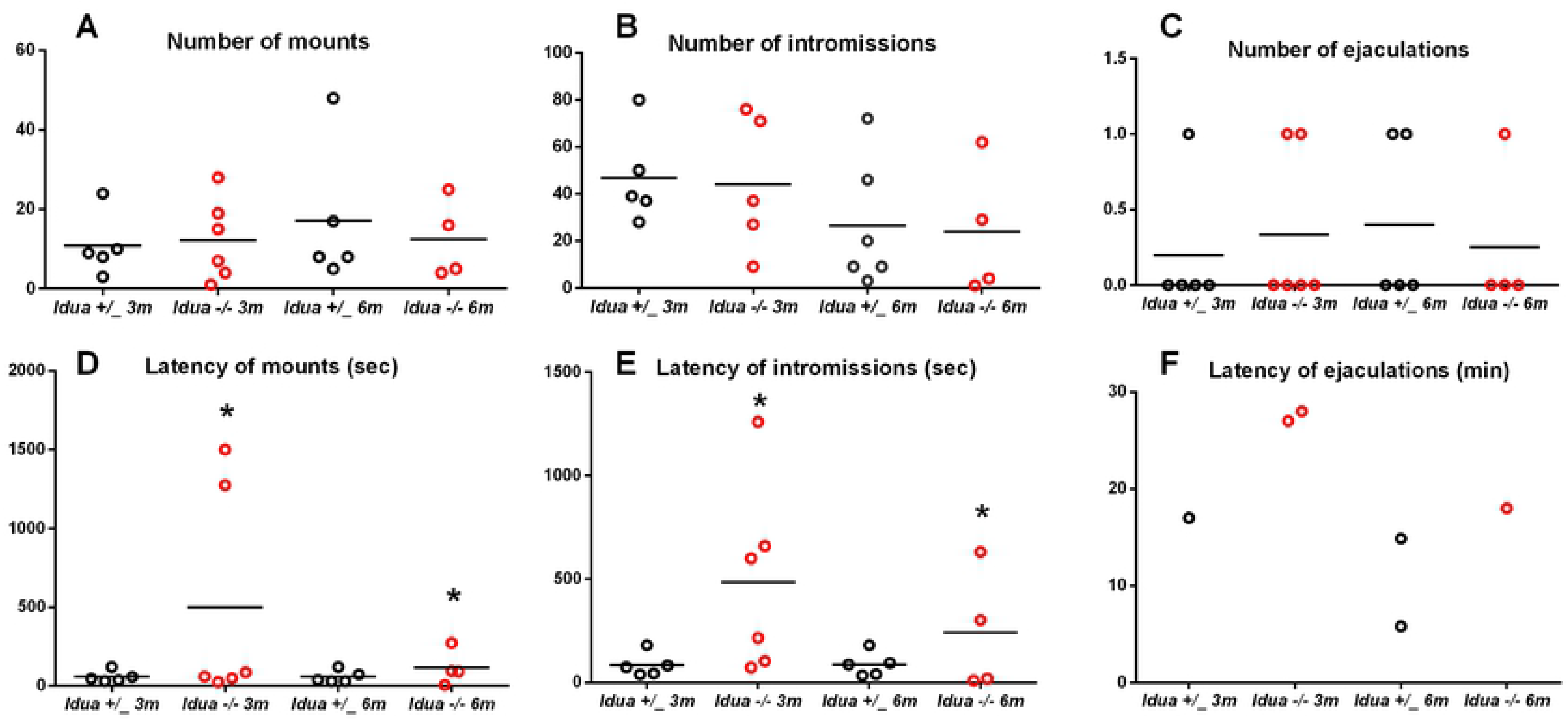
Analysis of male sexual behaviour. *Idua*+/_ and *Idua*−/− mice were previously adapted to mating and exposed to oestrous-induced females. In males, tests were performed in two different time points of MPS I progression: 3 and 6 months of age. The experiment was performed under dim-red light, for 30 minutes Statistical analysis: generalized estimating equations. *p<0.05 compared to control group.

From those mice whose reached ejaculation in 30 minutes, the number of ejaculatory episodes did not statistically differ between control and KO mice, independently of age (χ^2^ = 0.001, p = 0.971). Due to the low number of such copulatory behaviour, the latency of ejaculation could not be statistically compared between the groups (Fig 3F).

To explore the influence mobility issues may have exerted in these observed increased latency of mounts and intromissions, we performed new analyses for both variables including total quadrants crossed as a control variable. Number of quadrants had an effect on latency of mount (χ^2^ = 5.73, p = 0.017) but not latency of intromission (χ^2^ = 1.82, p = 0.18).

### Female sexual behaviour

There was no effect of genotype on both lordosis frequency (χ^2^ = 0.603, p = 0.44) or lordosis quotient (χ^2^ = 0.025, p =0.876) (Fig 4).

**Figure 4:**
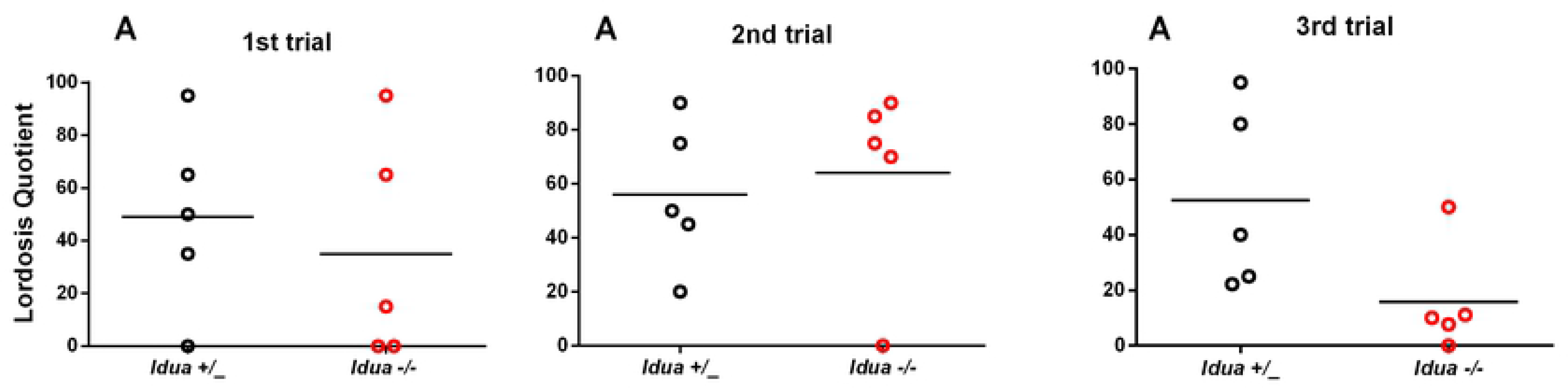
Analysis of female sexual behaviour. Two-month-old *Idua*+/_ and *Idua*−/− female mice were accompanied through three consecutive oestrous cycles. In the third proestrus phase, they were exposed to an experienced male to evaluate sexual behaviour. The analysis was repeated for the two following proestrus phases. The experiment was performed under dim-red light, for 15 minutes or until the completion of 20 mounts. Statistical analysis: generalized estimating equations.

### Hormonal plasmatic measurements

No effects of genotype were found on the plasmatic levels of oestradiol (χ^2^ = 0.74, p =0.39), progesterone (χ^2^ = 0.036, p = 0.85) or testosterone (χ^2^ = 0.82, p = 0.36) in female mice. Similarly, no effects of genotype were found on plasmatic levels of oestradiol, progesterone and testosterone in males (χ^2^ = 2.90 p= 0.088; χ^2^ = 0.12, p = 0.73; 1.47, p = 0.23, respectively) (Table 1).

**Table 1:** Analysis of plasmatic levels of steroid hormones. After behavioural analysis, both male and female mice were euthanized and their blood was collected for testosterone, progesterone and 17-β-oestradiol quantification.

|  | Male |  | Female |  |
| --- | --- | --- | --- | --- |
|  | *Idua* +/_ | *Idua* -/- | *Idua* +/_ | *Idua* -/- |
| Testosterone<br>(ng/dL) | 169,65 (101,27) | 226,21 (155,71) | 242.5 (104.0) | 186.79 (89.63) |
| Progesterone<br>(ng/dL) | 17684,10 (2777,85) | 18448,41 (491,15) | 37388.01 (46272.73) | 32449.08 (44732.45) |
| 17-β-stradiol<br>(ng/dL) | 61,74 (51,38) | 35,91 (4,53) | 1381.61 (2286.41) | 1788.55 (3836.23) |

Values are expressed by mean and standard deviation in parenthesis. Statistical analysis: generalized linear models.

## Discussion

This study aimed to elucidate the effects of IDUA deficiency on copulatory behaviour in the MPS I model. GAGs are important for reproductive physiology, and their distribution in the reproductive tract is closely associated with sexual hormonal levels [18–20]. While this is the first study to investigate sexual behaviour in an MPS I murine model, Soper el al. (1999) had previously provided a similar description in MPS VII mice [12]. In their study, reproductive performance of β-glucuronidase KO mice was increased after enzymatic replacement therapy, which suggests GAG storage may exert a negative influence in sexual behaviour. However, the authors attributed their results to the improvement in cognitive functions and mobility after treatment, since ovaries of treated females had follicles and corpora lutea and testes of treated males were capable of producing sperm [12].

In the present work we did not observe a significant impairment in copulatory behaviour of neither male nor female KO mice. While male MPS I mice had greater latencies of both mount and intromission, we attributed this to the impaired mobility observed in MPS I mice. However, after controlling for the influence of horizontal mobility, we concluded mobility may not influence intromission latency as much as mount latency. While courting behaviours, such as chasing the female through the testing arena, preclude the first mount and require great levels of agility [21], mobility may not be as important for performing the first instance of intromission after the first mount has been performed.

Control for copulatory behaviour is largely performed by the hypothalamic preoptic region [22]. MPS I mice have been shown to accumulate GAGs and secondary substrates, such as gangliosides and cholesterol, in their hippocampus and amygdala [23]. It is possible these substrates are also present in the hypothalamus, which could impair neuronal function and disturb regulation of copulatory behaviour.

Female KO mice crossed fewer total and central quadrants compared to their WT counterparts, which is in accordance with previously published findings [13]. Male KO mice, however, crossed more centre quadrants in the open field test, which is an unexpected behaviour in rodents since they prefer to walk near vertical surfaces [24]. This behaviour could evidence a cognitive impairment of KO mice.

Since steroidal hormones are synthesized from cholesterol and lysosome plays a crucial role in steroidogenesis [25], we suspected that disruptions in lysosomal hydrolases could impair hormone production. Testosterone, 17- β-estradiol and progesterone act on hypothalamic structures to control sexual appetite and behaviour [26–29]. However, in the evaluated time point of MPS I progression, levels of plasma testosterone in males and progesterone and 17- β-estradiol in females did not differ from their control counterparts, which could explain the normal pattern of copulatory behaviour observed in these mice.

The similarity of hormone levels between KO and control mice, in both genders, further suggest that higher latencies of male mounts and intromissions are related to motor dysfunction. It is possible we were unable to find differences in female copulatory behaviour between genotypes because male copulatory behaviour requires more mobility when compared to the female behaviour [30]. Thus, the motor limitation described in MPS I females appeared to not be relevant for sexual behaviour.

Ejaculation could not be observed in all male mice, independently of group and age. We attribute this result to the test duration, since mice may take longer than 30 minutes to reach the first ejaculation [15]. However, we adopted this test duration to avoid female pregnancies, since they were not ovariectomised and our main objective was to evaluate the male ability to execute the expected copulatory behaviours in rodents. It is important to note that two KO male mice presented a poor sexual performance during sexual behaviour test, however, both were able to fertilize healthy females during the habituation protocol of sexual activity [10]. This suggests reproductive performance overall is still successful in MPS I mice, despite the impairment in mobility observed in these animals. It is possible further improvements would be observed if mice were treated with enzyme replacement therapy, given this treatment has been shown to restore mobility in MPS I mice [31].

In conclusion, we found that IDUA deficiency does not impair copulatory behaviour in female and male mice. Moreover, this enzymatic deficiency also does not influence steroidal hormonal production.

## Aknowledgments

We thank the foundation support agencies: “Coordenação de Aperfeiçoamento de Pessoal de Nível Superior” (CAPES), “Conselho Nacional de Desenvolvimento Científico e Tecnológico” (CNPq; Fellowship to CCN, and VD’A) and the “Associação Fundo de Incentivo à Pesquisa” (AFIP). We also thank Dr. Monica Levy Andersen, who kindly donated us the synthetic hormones.

ABBM and CCN performed the experiments, analysed data and elaborated the manuscript, and VDA participated in the elaboration of study design and in the analysis of all data. All authors revised and contributed to the manuscript writing.

## Ethical approval

All applicable international, national, and/or institutional guidelines for the care and use of animals were followed.

## References

1. Neufeld EF, Muenzer J. The Mucopolysaccharidoses. In: Scriver CR, Sly WS, Childs B, Beaudet AL, Valle D, Kinzler KW VB, editors. The metabolic & molecular bases of inherited disease. New York: McGraw-Hill;2001. pp. 3421–3452.

2. Clarke LA. The mucopolysaccharidoses: a success of molecular medicine. Expert Rev Mol Med. 2008;1: 1–18.

3. Muenzer J, Wraith JE, Clarke LA. Mucopolysaccharidosis I : Management and treatment guidelines. Pediatrics. 2009;123: 12–29.

4. Kakkis ED. Enzyme-replacement therapy in mucopolysaccharidosis I. N Engl J Med. 2001;344(3): 182–188.

5. Hendriksz CJ, Moss GM, Wraith JE. Pregnancy in a patient with mucopolysaccharidosis type IH homozygous for the W402X mutation. J Inherit Metab Dis. 2004;27(5): 685–686.

6. Remérand G, Merlin E, Froissart R, Brugnon F, Kanold J, Janny L, et al. Four successful pregnancies in a patient with mucopolysaccharidosis type I treated by allogeneic bone marrow transplantation. J Inherit Metab Dis. 2009;Suppl 1: S111–S113.

7. Castorina M, Antuzzi D, Richards SM, Cox GF, Xue Y. Successful pregnancy and breastfeeding in a woman with mucopolysaccharidosis type I while receiving laronidase enzyme replacement therapy. Clin Exp Obstet Gynecol. 2015;42(1): 108–113.

8. Chung S, Ma X, Liu Y, Lee D, Tittiger M, Ponder KP. Effect of neonatal administration of a retroviral vector expressing α-l-iduronidase upon lysosomal storage in brain and other organs in mucopolysaccharidosis I mice. Molecular Genetics and Metabolism. 2007;90(2): 181–192.

9. Do Nascimento CC, Aguiar Junior O, D’ Almeida V. Analysis of male reproductive parameters in a murine model of mucopolysaccharidosis type I (MPS I). Int J Clin Exp Pathol. 2014;7(6): 3488–3497.

10. Do Nascimento CC, Aguiar O, Viana GM, D’Almeida V. Evidence that glycosaminoglycan storage and collagen deposition in the cauda epididymis does not impair sperm viability in the Mucopolysaccharidosis type I mouse model. Reproduction, Fertil. Dev. Forthcoming.

11. Milazzo J, Bironneau A, Vannier J, Liard-zmuda A., Macé B. Precocious initiation of spermatogenesis in a 19-month-old boy with Hurler syndrome. Basic Clin Androl. 2014 24(8): 1–7.

12. Soper BW, Pung AW, Vogler CA, Grubb JH, Sly WS, Barker JE. Enzyme replacement therapy improves reproductive performance in mucopolysaccharidosis type VII mice but does not prevent postnatal losses. Pediatric Research. 1999;45(2): 180–186.

13. Baldo G, Mayer FQ, Martinelli B, Dilda A, Meyer F, Ponder KP, et al. Evidence of a progressive motor dysfunction in Mucopolysaccharidosis type I mice. Behav Brain Res. 2012;233(1): 169–175.

14. Ohmi K, Greenberg DS, Rajavel KS, Ryazantsev S, Li HH, Neufeld EF. Activated microglia in cortex of mouse models of mucopolysaccharidoses I and IIIB. PNAS. 2003;100(4): 1902–1907.

15. Kikusui T. Analysis of Male Aggressive and Sexual Behavior in Mice. Methods Mol Biol. 2013;1068: 307–318.

16. Brock O, Baum MJ, Bakker J. The Development of Female Sexual Behavior Requires Prepubertal Estradiol. Journal of Neuroscience. 2011;31(15): 5574–5578.

17. Chinen CC, Faria RR, Frussa-filho R. Characterization of the Rapid-Onset Type of Behavioral Sensitization to Amphetamine in Mice: Role of Drug – Environment Conditioning, Neuropsychopharmacology. 2006;31: 151–159.

18. Sampaio LO, Markus, RP, Nader HB. The effect of sexual hormones on the sulfated glycosaminoglycan pattern of male genital accessory organs. Arch Biochem Biophys. 1985;240(1): 470–477.

19. Grunert G, Fernández S, Tchernitchin AN. Methods for the evaluation of responses to estrogen in individual cell types or regions of the uterus. Hormone Research. 1984;19(4): 253–262.

20. Sato E. Intraovarian control of selective follicular growth and induction of oocyte maturation in mammals. Proc Jpn Acad Ser B Phys Biol Sci. 2015;91(3): 76–91.

21. Hull EM, Wood RI, McKenna KE. Neurobiology of male sexual behavior. In: JD. Neill (Ed.), Physiology of Reproduction. New York: Elsevier;2006. pp. 1729–1824.

22. Wood RI, Newman SW. Integration of chemosensory and hormonal cues is essential for mating in the male Syrian hamster. J. Neurosci. 1995;15: 7261–7269.

23. McGlynn R, Dobrenis K, Walkley SU. Differential subcellular localization of cholesterol, gangliosides, and glycosaminoglycans in murine models of mucopolysaccharide storage disorders. J Comp Neurol. 2004;480(4): 415–426.

24. Lamprea MR, Cardenas FP, Setem J, Morato S. Thigmotactic responses in an open-field. Braz J Med Biol Res. 2008;41(2): 135–140.

25. Xie C, Richardson JA, Turley SD, Dietschy JM. Cholesterol substrate pools and steroid hormone levels are normal in the face of mutational inactivation of NPC1 protein. J Lipid Res. 2006;7(5): 953–963.

26. Durdiakova J, Ostatnikova D, Celec P. Testosterone and its metabolites modulators of brain functions. Acta Neurobiologiae Experimentalis (Wars). 2011;71(4): 434–454.

27. Arteaga-Silva M, Rodríguez-Dorantes M, Baig S, Morales-Montor J. Effects of castration and hormone replacement on male sexual behavior and pattern of expression in the brain of sex-steroid receptors in BALB/c AnN mice. Comp Biochem Physiol A Mol Integr Physiol. 2007;147(3): 607–615.

28. Hull EM, Domingues JM. Sexual behavior in male rodents. Horm behav. 2007;51(1): 45–55.

29. Matsumoto T, Honda S, Harada N. Alteration in sex-specific behaviors in male mice lacking the aromatase gene. Neuroendocrinology. 2003;77(6): 416–424.

30. Estep DQ, Lanier DL, Dewsbury DA. Copulatory behavior and nest building behavior of wild house mice (Mus musculus). Anim Learn Behav. 1975;3(4): 329–336.

31. Baldo G, Mayer FQ, Martinelli BZ, de Carvalho TG, Meyer FS, de Oliveira PG et al. Enzyme replacement therapy started at birth improves outcome in difficult-to-treat oorgans organs in mucopolysaccharidosis I mice. Mol Genet Metab. 2013;109(1): 33–40.

